# The Early Dodder Gets the Host: Decoding the Coiling Patterns of *Cuscuta campestris* with Automated Image Processing

**DOI:** 10.1101/2024.02.29.582789

**Authors:** Max Bentelspacher, Erik J. Amézquita, Supral Adhikari, Jaime Barros, So-Yon Park

## Abstract

*Cuscuta* spp., commonly known as dodders, are rootless and leafless stem parasitic plants. Upon germination, *Cuscuta* starts rotating immediately in a counterclockwise direction (circumnutation) to locate a host plant, creating a seamless vascular connection to steal water and nutrients from its host. In this study, our aim was to elucidate the dynamics of the coiling patterns of *Cuscuta*, which is an essential step for successful parasitism. Using time-lapse photography, we recorded the circumnutation and coiling movements of *C. campestris* at different inoculation times on non-living hosts. Subsequent image analyses were facilitated through an in-house Python-based image processing pipeline to detect coiling locations, angles, initiation and completion times, and duration of coiling stages in between. The study revealed that the coiling efficacy of *C. campestris* varied with the inoculation time of day, showing higher success and faster initiation in morning than in evening. These observations suggest that *Cuscuta*, despite lacking leaves and a developed chloroplast, can discern photoperiod changes, significantly determining its parasitic efficiency. The automated image analysis results confirmed the reliability of our Python pipeline by aligning closely with manual annotations. This study provides significant insights into the parasitic strategies of *C. campestris* and demonstrates the potential of integrating computational image analysis in plant biology for exploring complex plant behaviors. Furthermore, this method provides an efficient tool for investigating plant movement dynamics, laying the foundation for future studies on mitigating the economic impacts of parasitic plants.

## INTRODUCTION

*Cuscuta* species, commonly referred to as dodders, are obligate stem parasitic plants of the Convolvulaceae family, widely recognized as morning glories (Riviere, et al. 2013). *Cuscuta* infestations can have profound economic implications. It can infect and considerably reduce the yield of more than 25 different crop species and has been found in at least 55 countries across all continents (Kogan and Lanini 2005). *Cuscuta* infestations are difficult to control, and can reduce the yield of economically important plants such as tomato and alfalfa (Goldwasser, et al. 2012). *Cuscuta* is resistant to most of the commercially available herbicides (Nadler-Hassar and Rubin 2003), and a single plant can produce up to 15,000 seeds that can remain viable for at least 10 years (Saric-Krsmanovic and Vrbnicanin 2015). With environmental patterns being altered due to climate change, species distribution models predict that the field and host ranges of *Cuscuta* can expand even further (Cai, et al. 2022; Masanga, et al. 2021). Therefore, new control strategies focus on determining the molecular and environmental conditions that affect *Cuscuta*’s ability to detect and attach itself successfully to new hosts during its seedling stage (Hartenstein, et al. 2023). *Cuscuta* propagates through either seeds or stem cuttings (Ashton and Hutchison 1980; Hegenauer, et al. 2017). Following germination from seed or attachment to a host via stem cuttings, *Cuscuta* initiates a counterclockwise rotational movement, referred to as circumnutation. Circumnutation, first described by Charles Darwin (Darwin and Darwin 1880), is a phenomenon characterized by the combination of circular movement and axial growth for the independent and autonomous movements of young and growing organs such as the flowers, stems, shoots, tips, petioles, tendrils, or roots of plants, including *Cuscuta* (Agostinelli, et al. 2021; Brown 1993; Moulton, et al. 2020; Stolarz 2009; Wu, et al. 2020). Since this movement is crucial for *Cuscuta* to identify and secure a suitable host plant, circumnutation is the most essential process in early parasitism for the species. After recognition and successful attachment, *Cuscuta* starts producing feeding sites known as haustoria, which penetrate the vascular system of the host plant to obtain water and nutrients for its own survival (Jhu and Sinha 2022; Kim and Westwood 2015; Shimizu and Aoki 2019). Without attachment to a host, *Cuscuta* cannot survive independently due to its lack of leaves and roots. Unraveling these circumnutating, coiling, and attaching dynamics are key to understand how *Cuscuta* survives and interact with its environment.

Interestingly, circumnutation movement is influenced by the circadian clock (Niinuma, et al. 2005; Stolarz 2009). The circadian clock helps plants adapt to varying light conditions by regulating physiological and behavioral processes through an endogenous 24-hour rhythm (Creux and Harmer 2019; Greenham and McClung 2015; McClung 2006). In response to changing light conditions, the circadian clock uses external cues, such as light and temperature, to synchronize its internal rhythm with the external environment. Notably, plant circadian rhythms are self-sustaining (free running) and have an intrinsic, endogenous component that allows it to maintain a 24-hour cycle even in the absence of external cues (Stolarz 2009).

The conventional method for phenotyping and characterizing circumnutation and other developmental traits typically entails direct visual examination and annotation. This process is both time-consuming and prone to mistakes, which limits the number of phenotypes observed, and plant samples analyzed. In a digital data-dominated era, open-sourced high-throughput plant phenotyping (HTPP) pipelines are crucial to gain better and more profound insights from image-based data (Araus and Cairns 2014; Fahlgren, et al. 2015). For example, 2D image-processing HTTP pipelines can automatically track size changes in *Arabidopsis* leaves (Swartz, et al. 2023), detect plant-pathogen interactions in excavated maize roots (Pierz, et al. 2023), characterize leaf venation patterns for grasses (Robil, et al. 2021), and even trace circumnutation movements made by *Arabidopsis* stems (Mao, et al. 2023). However, adapting existing HTPP pipelines for *C. campestris* phenotyping presents challenges due to several unique features of the organism. These include its absence of leaves and root system, as well its highly variable movement and twinning patterns.

In this study, we introduce a Python-based image processing pipeline designed to phenotype key traits associated with *C. campestris* circumnutation and coiling. This pipeline allows the automated detection of coiling location, angle, initiation, and completion times, along with the duration of coiling stages in between. Subsequently, we applied the pipeline to gain a better understanding of circumnutation and coiling of *C. campestris* and investigated how circadian rhythm influences successful inoculation. The findings obtained from this automated image analysis tool aligned with those derived by manual annotation, confirming the reliability of the Python pipeline. The pipeline is openly accessible as a series of commented Jupyter notebooks to encourage appropriate modifications for different image data or experimental setups. This approach not only enhances the precision in tracking plant movements but also paves the way for future research on the mechanisms underlying parasitic plant interactions and potential sustainable weed control strategies.

## MATERIALS AND METHODS

### Plant growth and *C. campestris* inoculation

The experimental workflow of our study is illustrated in Figure 1. *C. campestris* seedlings were inoculated on one-month-old *Beta vulgaris* in a greenhouse with a temperature of 25-30°C and day/night cycle of 16/8 hours. Three weeks after the initial inoculation, mature *C. campestris* stem segments were collected from the greenhouse. To evaluate the potential for inducing coiling under our experimental conditions, preliminary tests were conducted by attaching these *C. campestris* stems to 3-week-old *Arabidopsis* Col-0 and to bamboo skewers (30 cm in length and 4 mm in diameter), enabling a comparison of coiling patterns on living versus non-living materials (Fig. 2a-b, Suppl. Video 1). The *C. campestris* stem segments used in the inoculation experiments measured between 6-8 cm, with shoot tips ranging from 4-6 cm in length and a single offshoot branch. The *C. campestris* stems were attached to their host or skewer with pieces of 3M Magic tape of approximately 3-3.5 cm in length and 4 mm in width. The tape was applied on the shoot tip at half its length. To ensure optimal inoculation, the shoot tips were aligned facing left at an angle of 30-60 degrees, considering that *C. campestris* shoot tips exhibit an anti-clockwise movement pattern. The attached shoot tips were kept under 24-hour constant lightening using far-red enriched lights with an intensity of 100 µmol/m^2^ (GE R30 Soft White Medium Base, model #30711) as previously described in (Bernal-Galeano, et al. 2022) (Supp Fig. 1). For the time lapse experiment, every sampling setup consisted of five *C. campestris* stems attached to five skewers roughly equidistant from each other. This setup was repeated seven times for three different inoculation times: 9 AM, 12 PM, and 4 PM, resulting in 35 individual stems for every inoculation time.

**Fig 1.**
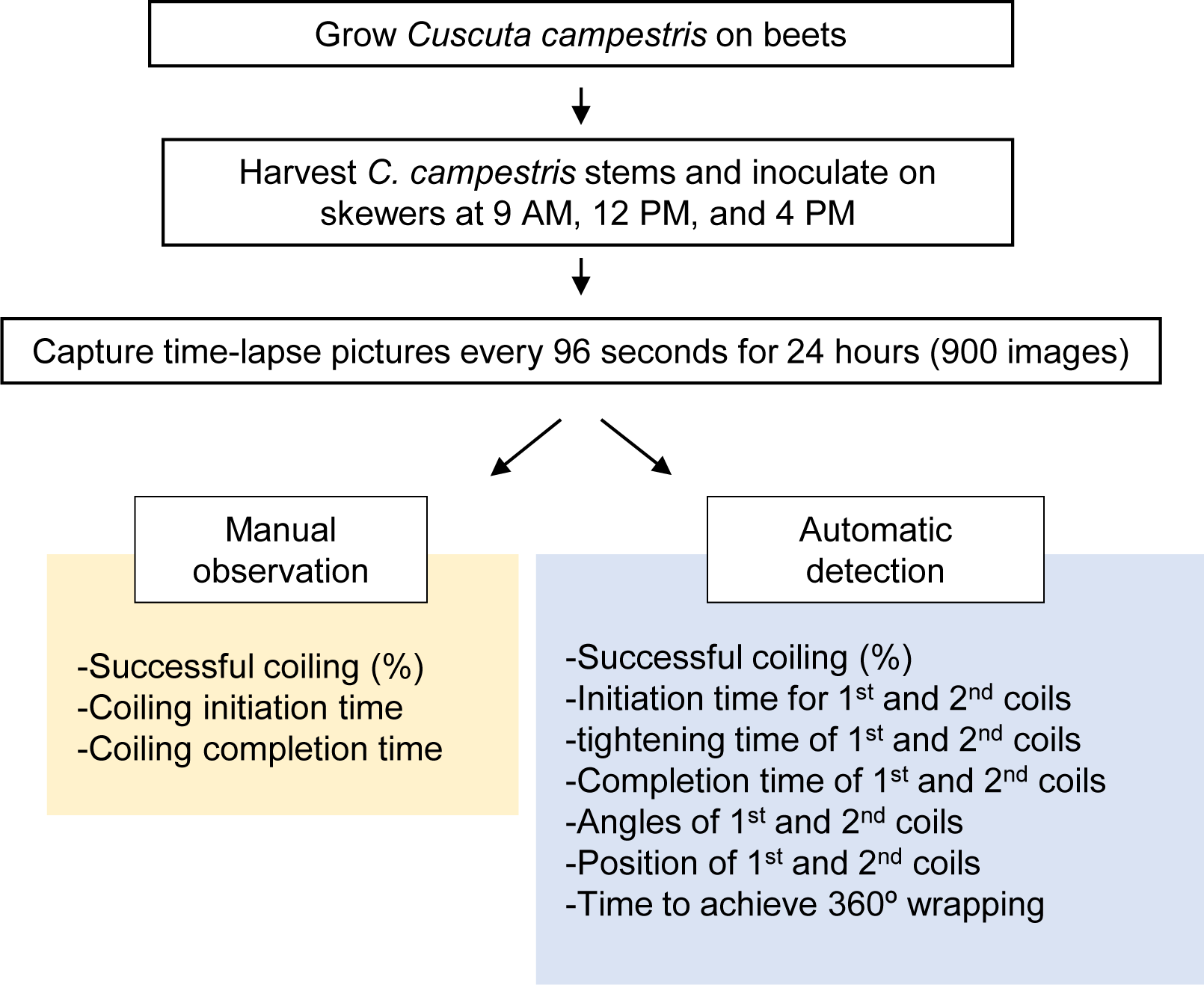
Experimental workflow for monitoring the coiling movement of *Cuscuta*. After acquiring time-lapse images of *Cuscuta* stems subjected to varying inoculation times, each measurement was analyzed through manual observation or automatic detection.

**Fig 2.**
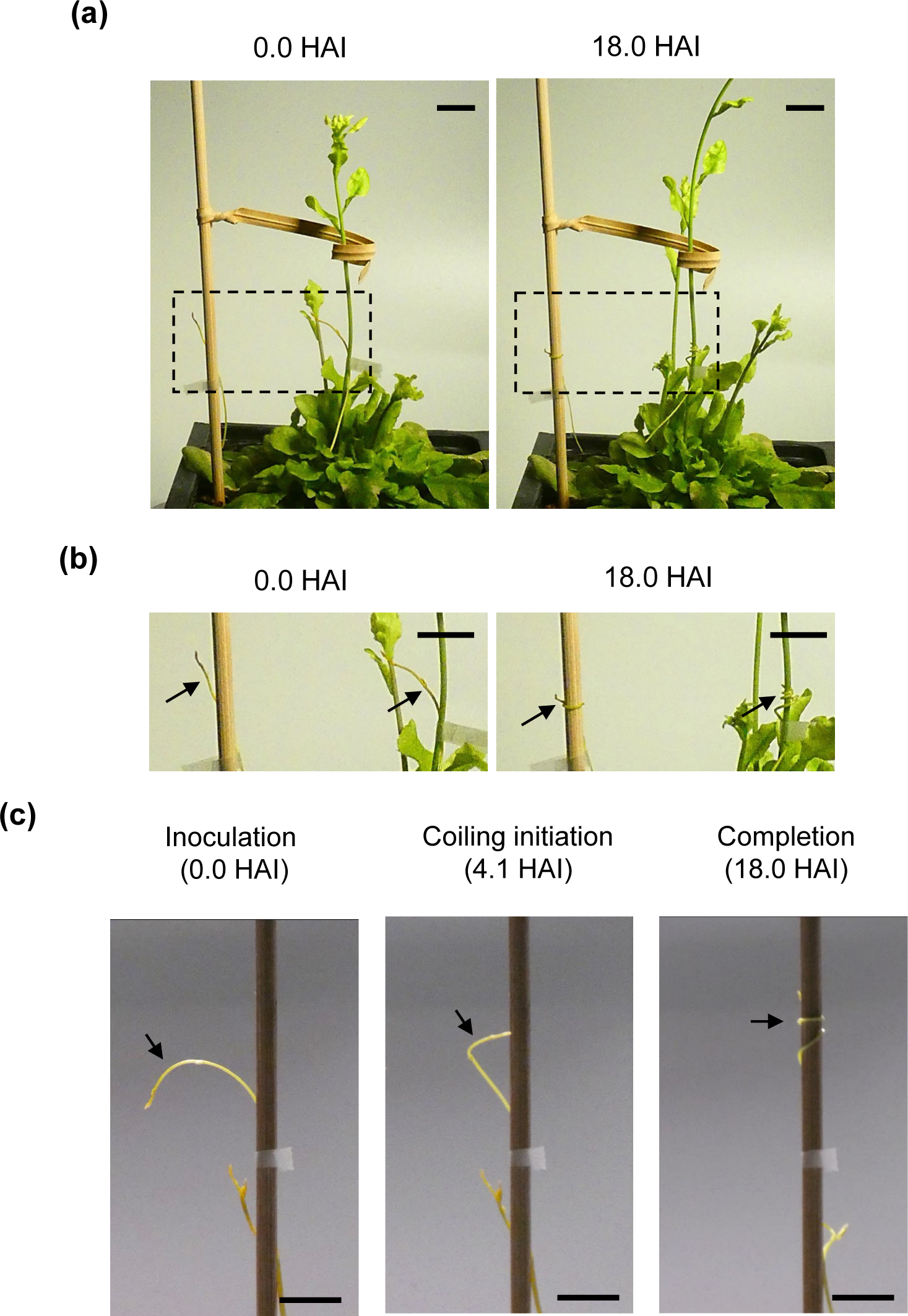
*Cuscuta* stems were wrapped around *Arabidopsis* stems and bamboo skewers. **(a)** *Cuscuta* stems were attached on *Arabidopsis* Col-0 and bamboo skewers and incubated for 24 hours. **(b)** Zoomed-in images of lined boxes in **(a)**. **(c)** Timelapse images were captured to observe the coiling movement of *Cuscuta* stems under different inoculation times. Scale bar: 1 cm. HAI: hours after inoculation. Black arrows indicate *Cuscuta* stems.

### Camera setting, time lapse, and videos

Photos were captured on a Panasonic DC-FZ80 digital camera. The aspect ratio was set to 16:9, picture quality to 4896 x 2752 pixels, aperture value of 4.0, shutter speed of 100, ISO 1600, and an interval time of 96 seconds. The white balance was set manually based on the light conditions in the room before the timelapse. The focus was set using automatic settings, then switched to manual prior to capture. Flash, image stabilizing, and automatic white balance settings were turned off. From the ruler attached next to the skewers, 14 pixels correspond to 1 mm length. Shotcut software (https://shotcut.org) was used to make and render timelapse videos with the image sequence option. The videos were rendered at 30fps using a libx265 HEVC codec and exported as an mp4 file (Suppl. Videos 2-4).

### Manual observation

According to *C. campestris* movement and coiling patterns on the bamboo skewer, we manually observed success rate of coiling, coiling initiation, and completion times while *C. campestris* stems coiled on bamboo skewers. Successful inoculations were quantified by evaluating whether the *C. campestris* stems exhibited complete coiling around the bamboo skewers showing a stable physical interaction. Only *C. campestris* stems that successfully coiled on bamboo skewers were considered for analysis. Coiling *initiation* was defined as the time from the point of contact until the *C. campestris* stems began its characteristic anti-clockwise twining motion. *Completion* of inoculation was determined when the main shoots ceased to move significantly around the surrogate host (Fig. 2c). Time intervals for initiation and completion of coiling were recorded and averaged in a spreadsheet for each inoculation event within the experiment.

### Image processing setup

An in-house SciPy-based Python image processing script was developed to automatically extract movement and position information from the individual standstill photos described above. (Suppl. Videos 5-9). This way we divided the *C. campestris* coiling process into nine partially overlapping steps: 1) inoculation, 2) Coil 1 initiation, 3) Coil 1 positioning, 4) Coil 1 tightening, 5) Coil 1 completion, 6) Coil 2 initiation, 7) Coil 2 positioning, 8) Coil 2 tightening, and 9) completion, as explained below. The script is based on a combination of color, hue, and saturation thresholding and elementary image morphology operations. All the image processing and detailed analyses steps, limitations, and considerations can be read as Jupyter notebooks available at the GitHub repository listed at the end. The exact hyperparameters were first manually tuned for a single photo and these values were later used for the rest of the images; this was possible since all the photos have comparable illumination and color values.

### Automated image cleaning and annotation

First, the background was thresholded (black in Fig. 3c); the centers of the skewers were approximated with ordinary least-squares best fit lines (red in Fig. 3c); and the top edge of the tape pieces was located (orange in Fig. 3c). Next, the image was split into 5 overlapping sub-images, each centered on a different skewer; left and right limits determined by neighboring skewers; bottom margin determined by tape (Fig. 3d-h). The *C. campestris* in each section was identified as the union of large, connected components near the central skewer (Fig. 3i-m). The resulting segmented *C. campestris* was skeletonized (Zhang and Suen 1984) (gray in Fig. 3n-r). Subsequent phenotyping was limited to instances where the skeleton crossed in front the center of the skewer (white dot in Fig. 3s-w). These crossing instances were modeled by weighted linear RANSACs (Fischler and Bolles 1981), where pixels closer to the crossing site had higher weights (light blue and mustard in Fig. 3t-w). The crossing angle was determined as the complement of the angle between the skewer central line and the RANSAC line (pink in Fig. 3t-w). It is important to note that an angle of 0 degrees indicates that the *C. campestris* crosses perpendicularly to the skewer. Repeating these steps for all snapshots resulted in location and angle time series for Coil 1 and Coil 2 for every individual *C. campestris* (Fig. 4a). Missing time points were interpolated using linear splines and outliers were removed following a Savitzky-Golay filter (Savitzky and Golay 1964). A *C. campestris* stem was determined as successfully coiled if it presented relevant angle and position information for both Coil 1 and Coil 2.

**Fig. 3.**
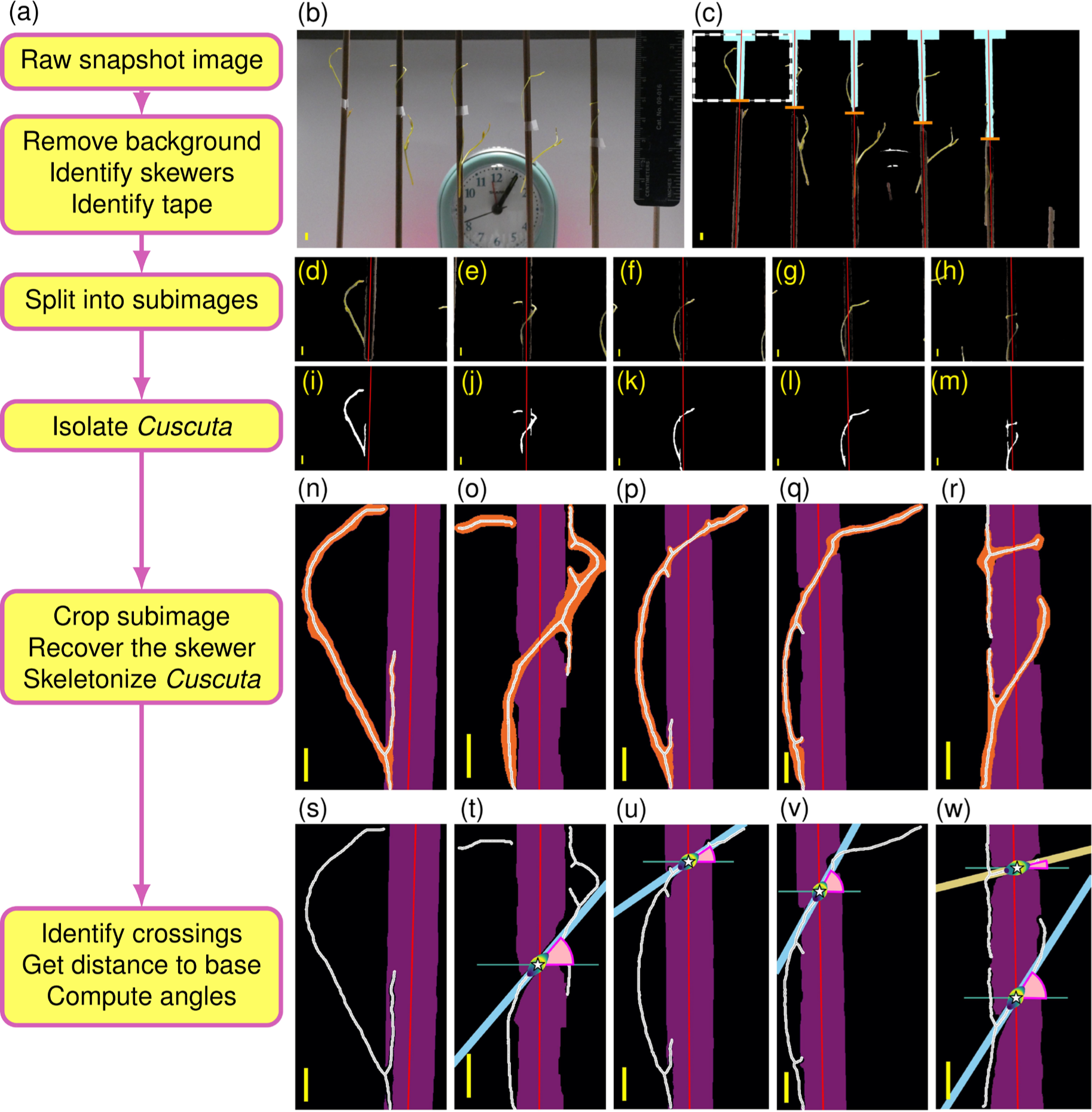
Image processing and analysis of *Cuscuta* images. **(a)** Summary of image processing steps performed, starting from a raw image and finishing annotations for each *Cuscuta*. Examples of each step are to the left. **(b)** Standstill image of the *Cuscuta* experiment setup as initial input. **(c)** The background was thresholded out (black), the center of each skewer was approximated with a best-fit line (red line), and the tape upper border was identified (orange line). Further steps only considered the skewer section above such tape (cyan). **(d-h)** The image was then split into 5 sub-images, each centered at a different skewer, from which **(i-m)** the relevant *Cuscuta* (white) was segmented and **(n-r)** skeletonized (gray). Further analysis focused solely on instances where the skeleton crosses the center of the skewer. **(s)** Example where no information is extracted as the *Cuscuta* has not yet passed in front of the skewer. **(t-w)** The position where *Cuscuta*’s skeleton crosses the skewer is recorded (white star) as well as the complementary angle (pink pie) made by the first coil (cyan line) and **(w)** second coil (mustard line). The skewer is shown in purple and its midline approximation is shown in red. The yellow scalebar corresponds to 56 pixels which approximates to 2 mm for all images.

**Fig. 4.**
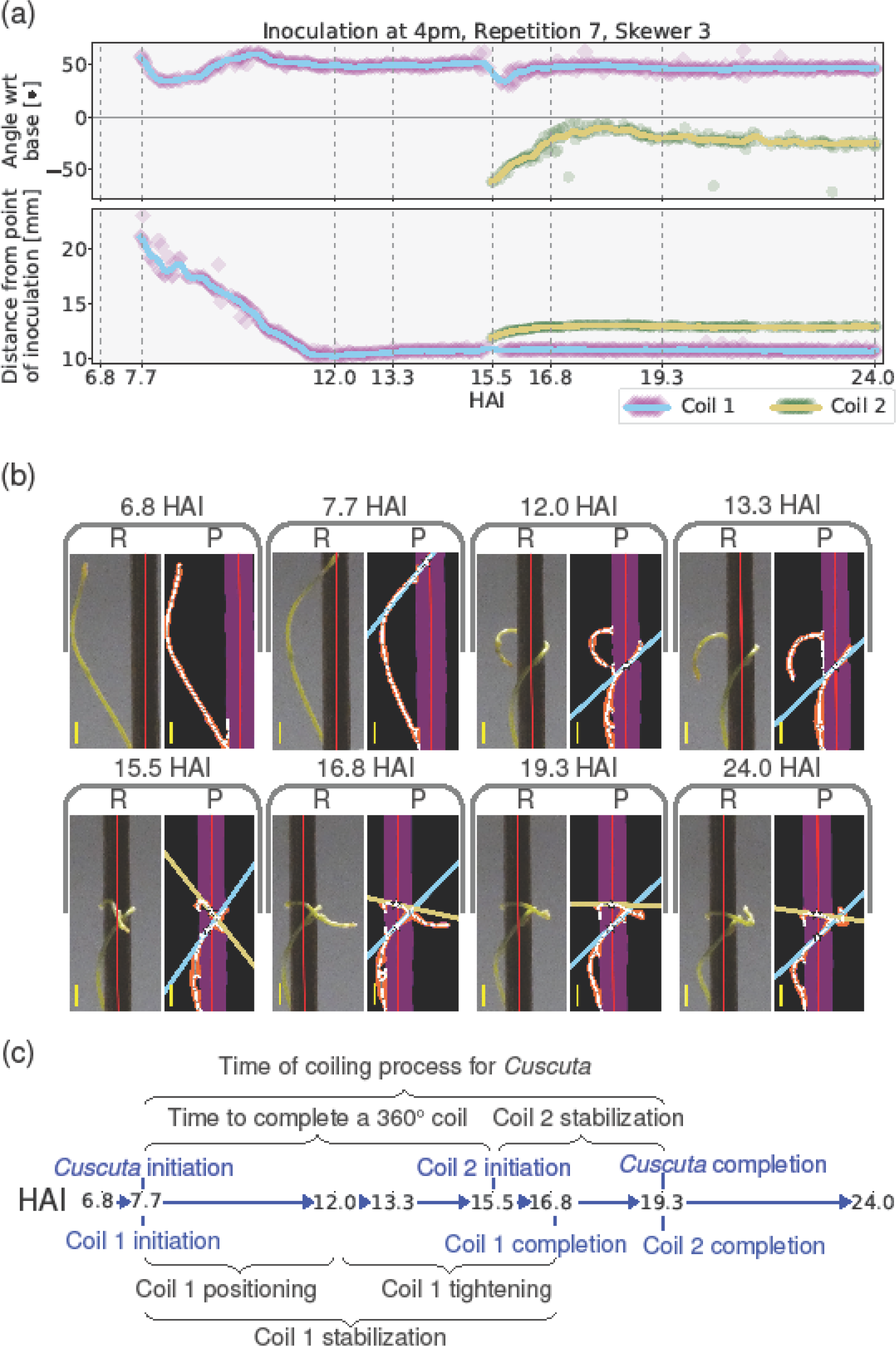
Automated detection of essential time points of *Cuscuta* coiling process. **(a**) For a single *Cuscuta,* the angle and position information of both coils is extracted from all the standstill images (Fig. 3). The scatter plot represents the actual observations. The solid lines represent the time series obtained after removing outliers and smoothening the signal. **(b)** Sample standstill sub-images represented by the time series. Raw (R) and its corresponding processed (P) sub-image are presented side by side for visual comparison. The yellow scalebar corresponds to 2 mm for all sub-images. In this analysis, the time series contains no information prior to 7.7 HAI since *Cuscuta* has not crossed in front of the skewer yet. Similarly, no information on Coil 2 was recorded prior to 15.5 HAI. Refer to Supplementary Video 5 for an animated example. **(c)** From the time series, for each coil we can immediately determine the time points when it initiated, when its position stopped changing, and when its angle stopped changing. By identifying these timepoints (in blue), we can deduce the duration of coiling stages (in gray).

### Automated feature extraction

As illustrated in Figure 4, Coil 1 is *initiated* at 7.7 hours after inoculation (HAI) when the *C. campestris* crosses in front of the skewer for the first time. Immediately after, Coil 1 *positions* itself for the next 4.3 hours—both its angle and position vary more than 5 degrees and 1 mm respectively with respect to their final values at 24 HAI. In this case, Coil 1 stops positioning at 12.0 HAI, as its position remains stable for the remainder of the recording. Coil 1 then *tightens* for the next 4.8 hours —its angle remains variable while its position is stable. Finally, at 16.8 HAI, Coil 1 is *completed* as its angle also remains stable for the remainder of the recording. The coil *stabilization* time is simply the difference between completion and initiation times, which is 9.1 hours in this case. Note that this is the same as the sum of positioning and tightening periods. The times and periods for Coil 2 are analogous. In this case, Coil 2 initiates at 15.5 HAI, it exhibits no positioning period, and it tightens for 3.8 hours to be completed at 19.3 HAI so that it is stabilized after 3.8 hours. Additionally, the time for *C. campestris* to turn 360° was determined as the difference between Coil 1 and Coil 2 initiation times (7.8 hours). Finally, the whole coiling process time for a *C. campestris* was determined as the time difference between the initiation of Coil 1 and the completion of Coil 2 (11.6 hours). An animated version of this example can be seen in Supplementary Video 5.

### Statistical analysis

A Mann-Whitney U-test (Mann and Whitney 1947) was performed to compare if the reported position, angle, and time values had statistically different average values for different pairs of initial inoculation times or different coils. This test was chosen since it does not assume any special conditions on the underlying distributions. This test was used whenever comparing manual or automatic observations.

Spearman’s rank correlation coefficients were computed between the time to turn 360° and the different time periods related to the coiling —positioning, tightening, stabilization times— as well as between positioning and tightening times. Additionally, p-values associated to these coefficients were computed following a t-distribution. These analyses were carried out separately for different inoculation times and for Coil 1 and Coil 2. (Supp Fig 4). The Spearman coefficient was chosen because it does not assume linearity.

## RESULTS

### *C. campestris* coiled on both living and non-living hosts in 24 hours

Within 24 hours following *C. campestris* inoculation, the shoot tips successfully coiled on both bamboo skewers and *Arabidopsis* plants (Fig. 2a-b and Suppl. Video 1). This indicates that *C. campestris* can coil around a host within a day, and that bamboo skewers are suitable surrogate hosts for studying *C. campestris* coiling movements. These findings are consistent with previous research showing *C. campestris*’s ability to coil around non-living materials and form haustoria when exposed to an appropriate blue/far-red light ratio (Bernal-Galeano, et al. 2022; Hegenauer, et al. 2017). Consequently, skewers were used as surrogate hosts for subsequent analysis. Furthermore, we conducted manual observations to assess the success rate of coiling, as well as to track the initiation and completion times of coiling, while closely monitoring *C. campestris* stems as they coiled around bamboo skewers (Fig 2c).

Success rate of *C. campestris* coiling under different inoculation times

Manual observations indicated a 100% success rate for *C. campestris* coiling at 9 AM (35 stems), whereas the success rates were approximately 83% at both 12 PM and 4 PM, with 29 stems each (Fig 5a, left panel; Supplementary Table 1, and Suppl. Video 2-4). Automated observations showed a similar trend with lower rates for each time point: 97.1% at 9 AM (34 stems), 71.4% at 12 PM (25 stems), and 80% at 4 PM (28 stems) (Fig 5b, right panel; Supp table 2). This discrepancy is primarily attributed to difficulties in distinguishing the color between the *C. campestris* stem and the skewers in some replicates and in extracting data from skewers that did not remain completely stationary.

**Fig 5.**
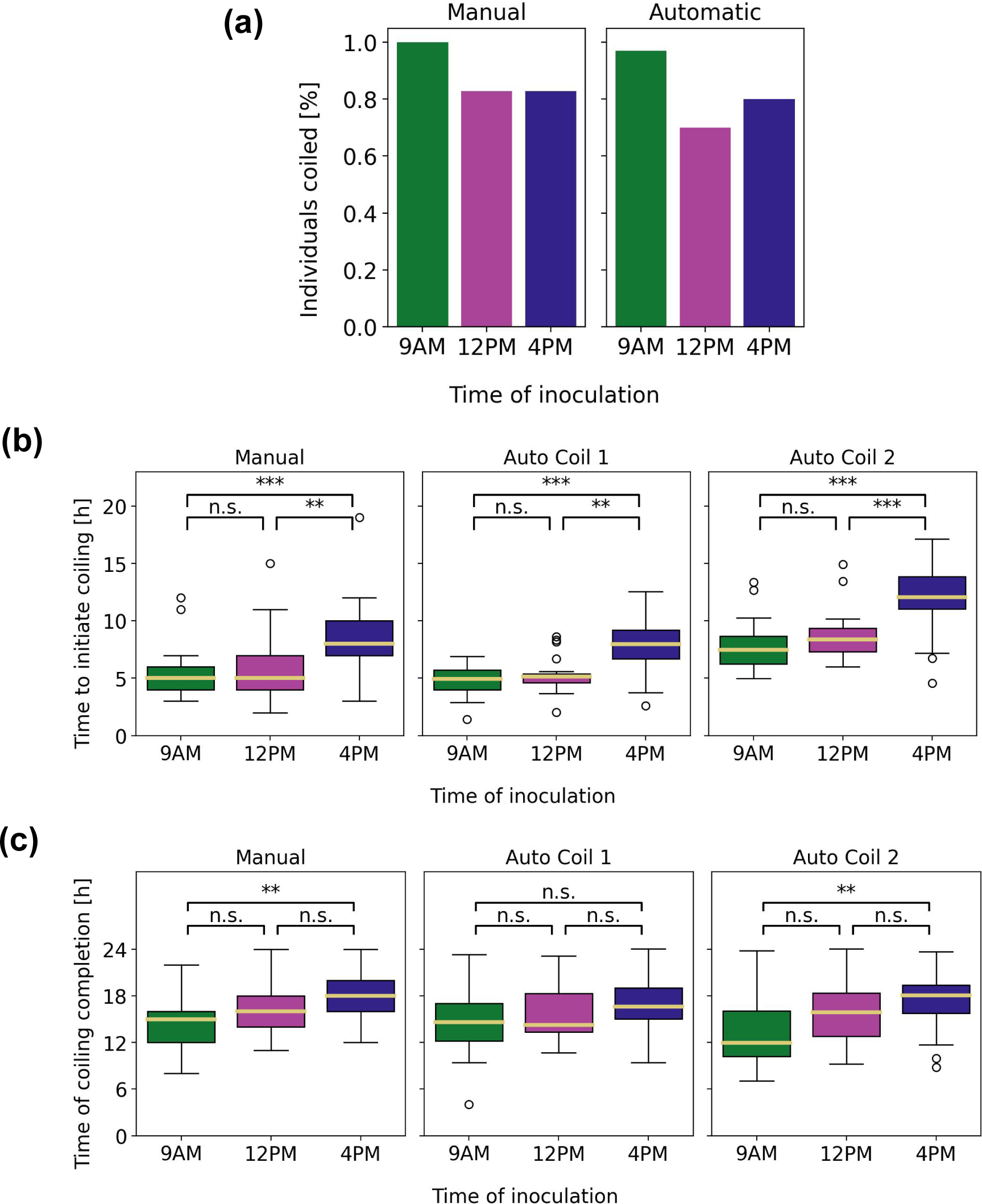
Examining the coiling behavior of *Cuscuta* through manual and automatic detection methods. The successful rate of *Cuscuta* coiling (n=35) **(a)**, coiling initiation time **(b)**, and coiling completion time **(c)** were measured by manual and automatic detection methods. ns: Not significant, p-value of ≥ 0.05; *: p-value of 0.05 to 0.005, **: p-value of 0.005 to 0.0005, and ***: p-value of < 0.0005.

### Initiation and completion of *C. campestris* coiling under different inoculation times

Manual observations revealed average coiling initiation times of 5.3, 5.9, and 8.5 hours after inoculation (HAI) for *C. campestris* inoculated at 9 AM, 12 PM, and 4 PM, respectively (Fig 5b, left panel; Supp table 3, and Suppl. Video 2-4). Notably, significant differences were observed in initiation times between *C. campestris* inoculated at 9 AM and 4 PM (p-value < 10^-5^), as well as between 12 PM and 4 PM (p-value < 10^-3^). Interestingly, stems inoculated at 4 PM exhibited delayed coiling initiation compared to those at 9 AM, potentially due to longer resting periods or circumnutation before recognizing the bamboo skewer. However, no significant differences were observed between inoculation at 9 AM and 12 PM. A similar trend was observed in the completion times, with an average time of 14.4, 16.5, and 18.0 HAI for 9 AM, 12 PM, and 4 PM, respectively (Fig 5c, left panel; Supp table 3, and Suppl. Video 2-4). In this case, the completion time was significantly different between *C. campestris* inoculated at 9 AM vs 4 PM (p-value < 10^-3^).

Similar conclusions can be drawn from the automatic observations. Coil 1 was initiated on average at 4.8, 5.3, and 7.9 HAI for 9 AM, 12 PM, and 4 PM, respectively (Fig 5b, middle panel; Supp table 4). Coil 2 was initiated at 7.8, 8.7, and 11.9 HAI for 9 AM, 12 PM, and 4 PM, respectively (Fig 5b, right panel; Supp table 5). These times are significantly different between *C. campestris* inoculated at 9 AM and 4 PM (p-values < 10^-5^), and between 12 PM and 4 PM (p-values < 10^-4^) for both Coil 1 and Coil 2. There were no significant differences between 9 AM and 12 PM. Do notice that the automated observations are based on when the *C. campestris* crossed in front of the skewer without regard the context of the movement, which most likely explains why the automatically reported times are earlier than the manual ones.

The automatic observations show that Coil 1 was completed on average at 14.8, 15.9, and 16.7 HAI for 9 AM, 12 PM, and 4 PM, respectively (Fig 5c, middle panel; Supp table 4). Coil 2 was completed at 13.3, 15.2, and 17.4 HAI for 9 AM, 12 PM, and 4 PM, respectively (Fig 5c, right panel; Supp table 5). We observed that Coil 2 tends to reach a stable state slightly earlier than Coil 1, but these differences were not significant. There are no significant differences in completion time for Coil 1 with respect to inoculation time. However, there is a significant difference in Coil 2 completion times between 9 AM and 4 PM (p-value < 10^-3^). Since the automatic annotation focuses solely on the behavior of the *C. campestris* in front of the skewer, it ignores the possibility of the *C. campestris*’s upper tip still moving while the coil in front of the skewer is stationary. This could explain why the automatically reported completion times of Coil 2 happen earlier than the manual ones. Nonetheless, despite different annotating criteria, both manual and automatic observations highlight significant differences in coiling initiation and completion times among *C. campestris* inoculated at different times, with those inoculated at 9 AM consistently showing earlier coiling compared to those inoculated at 4 PM.

### The automated pipeline tracks the coiling stabilization, twisting times, angles, positions, and gaps of the *C. campestris* coils

The automatic observations revealed consistent findings across different inoculation times. Firstly, there were no significant variations in the gap size between coils or in the time required to complete a full 360° coil around the skewer (Supp Fig 2a-b). Additionally, examination of Coil 1 and Coil 2 angles at 24 HAI showed no statistical differences in their final angle and position across all inoculation times. The average angle of Coil 1 was 23.5, 21.8, and 30.4 degrees for *C. campestris* inoculated at 9 AM, 12 PM, and 4 PM, respectively (Supp Fig 2c, left panel), while for Coil 2, the angles were 8.1, 9.0, and 13.9 degrees, respectively (Supp Fig 2c, right panel). Notably, there were no statistical differences in final angles and positions observed across all inoculation times, whether examining Coil 1 or Coil 2 (Supp Fig 2c-d). However, a marked contrast was observed between the final angle of Coil 1 and Coil 2 for every inoculation time (p-values < 10^-4^) (Supp Fig 2c). Specifically, Coil 1 tended to be more angled or slanted, while Coil 2 generally aligned parallel to the ground. These results indicate that the coiling dynamics of *C. campestris*, including the speed of wrapping and the final positioning of the coils, are not influenced by the time of day at which inoculation occurs, but they might be affected by the mechanical differences between the first and second coils.

Similar trends were observed when examining the time periods of different coiling stages. There were no significant differences in the positioning, tightening, and stabilization period durations for either Coil 1 or Coil 2 across all inoculation times (Supp Fig 3a-c). However, there was a striking statistical difference between the positioning duration of Coil 1 and Coil 2 across all inoculation times (p-values < 10^-4^). Coil 1 tended to position for about 6.0, 5.6, and 5.6 hours, while Coil 2 only positioned for 1.4, 2.6, and 2.0 hours, for *C. campestris* inoculated at 9 AM, 12 PM, and 4 PM, respectively. These time differences between coils are not observed in tightening time duration, but they reemerge when considering stabilization times (p-values < 10^-3^). In this case, Coil 1 tends to stabilize within 10.0, 10.6, and 9.0 hours compared to 5.6, 7.0, and 5.4 hours for Coil 2 for inoculations at 9 AM, 12 PM, and 4 PM, respectively. In other words, Coil 1 tends to move up and down the skewer for a while before finding a stationary position, while Coil 2 remains in place soon after it has twined with just slight adjustments in angle, suggesting mechanical differences between these two coils (Supp Fig 3a-c).

There were no significant differences when comparing the whole duration of the coiling process across all inoculation times. This conclusion is supported whenever using either the manual or automated annotation criteria (Supp Fig 3d). Such result suggests that while *C. campestris* inoculated at 9 AM tends to initiate coiling considerably earlier than *C. campestris* inoculated at 4 PM, once they start, the mechanistic coiling processes and stages have similar durations. This could indicate that *C. campestris* inoculated at 9 AM reaches a stationary state considerably earlier than *C. campestris* inoculated at 4 PM mainly because the former has an earlier head start (Fig 6). This might also indicate that the mechanisms involved in coiling initiation are independent from those involved in turning and twining. This final observation is also supported by a lack of correlation between the time it takes *C. campestris* to complete a 360° turn and the time it spends positioning, tightening, or stabilizing, regardless of the inoculation time or the coil observed (Supp Fig 4).

**Fig 6.**
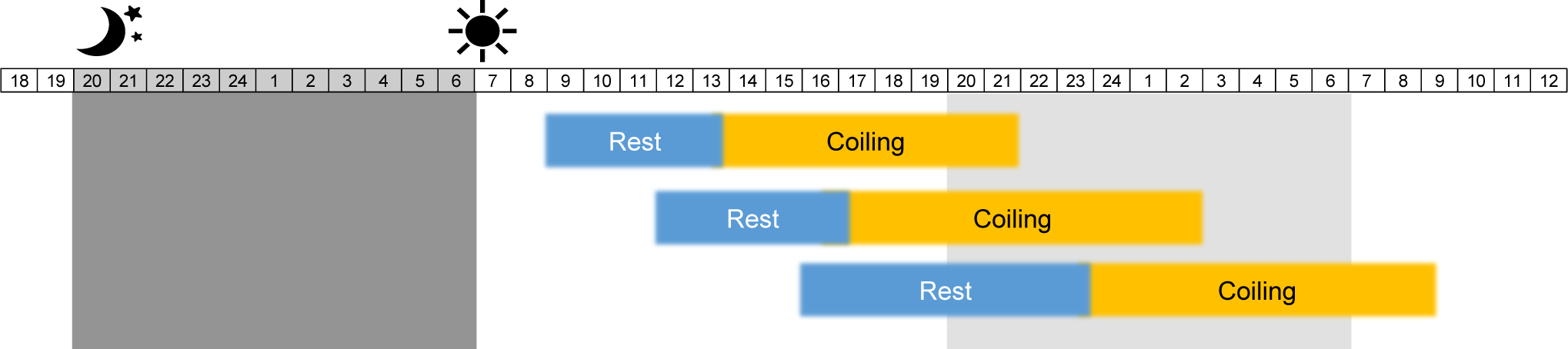
A summary of *Cuscuta* coiling movement using automatic detection methods with three different inoculation times (9 AM, 12 PM and 4 PM). Blue bars and yellow bars indicate average time of rest and coiling period. Dark and light gray shading indicates actual light/dark cycle and subjective darkness, respectively.

## DISCUSSION

Throughout evolution, plants have developed a plethora of survival strategies. Among these, parasitic plants have evolved unique adaptations to exploit other plant species for their resources. These unique members of the plant kingdom exhibit either facultative or obligatory dependence on other plant species for survival. They are also classified either as stem or root parasites, depending on the point of attachment to a host (Těšitel 2016). *Cuscuta* has evolved specialized haustoria structures and exhibits remarkable mobility, while undergoing atrophy in its root and leaf organs (Jhu and Sinha 2022; Sun, et al. 2018). Despite the critical role of mobility, such as circumnutation and coiling, in the successful parasitism of *Cuscuta*, studies investigating this aspect have been limited and primarily focused on manual observations, often in the context of responses to volatiles or light wavelengths (Furuhashi, et al. 2021; Furuhashi, et al. 2011; Runyon, et al. 2006; Yokoyama, et al. 2023). Addressing this gap, our study aimed to provide a combined analysis of *Cuscuta* coiling dynamics, using both manual observations and automated detection methods.

We found that *C. campestris* inoculated at 9 AM resulted in the highest success rate of inoculation and showed the fastest coiling initiation around the host (Fig. 5a-b). In contrast, inoculations at 4PM showed a lower success rate and an approximately 1.5-fold slower coiling initiation compared to those at 9 AM and 12 PM, possibly due to an extended resting stage or prolonged circumnutation movement (Fig 5b and 6). These observations suggest a difference in circumnutation and coiling behavior of *C. campestris* depending on the time of day. These observations give evidence that *C. campestris* perform circumnutation best in morning, moderately in afternoon, and lower in the evening, indicating that *C. campestris* has preferable times for active circumnutation and subsequent coiling. Interestingly, similar diurnal patterns have been observed in sunflowers, where young sunflowers exhibited a wide amplitude of circumnutation from midnight to noon, gradually decreasing in the afternoon and reaching a resting stage from evening (around 6PM) until midnight (Stolarz 2009). This indicates that like sunflowers, rootless and leafless *C. campestris* retains the light/dark period even though its stem is cut. Moreover, it suggests that *C. campestris* has its own internal oscillator capable of detecting the circadian clock, regulating circumnutation for successful parasitism independent of host-derived hormones or signals. According to previous reports, climbing plants have been reported to perform circumnutation more regularly (Baillaud 1962; Millet, et al. 1984; Yoshihara and Iino 2005). Since *Cuscuta* belongs to the morning glory family (Convolvulaceae), it is expected that *Cuscuta* will also perform circumnutation like other climbing plants. Additionally, it has been reported that *Early flowering 3* (*ELF3*) or *Timing of cab expression 1* (*TOC1*) regulates circumnutation (Niinuma, et al. 2007; Niinuma, et al. 2005), but in *Cuscuta australis*, it has been reported that not only *ELF3/4* but also *Response regulator 3 and 4* (*ARR3/4*)*, Cycling Dof factor 1 and 3* (*CDF1/3*)*, Flavin-binding, kelch repeat 1* (*FKF1*), and *Cryptochrome-interacting basic-helix-loop-helix 1* (*CIB1*) are missing (Sun, et al. 2018). It suggests a research gap in understanding how *Cuscuta* regulates its circumnutation despite lacking some key circadian clock and photoperiod genes. Therefore, although *Cuscuta* is influenced by existing circumnutation mechanisms like climbing plants, how it performs circumnutation well without these circadian clock genes including *ELF3* is a subject that needs to be researched in the future. In addition, it has been observed that the concentration of cytoplasmic free calcium (Ca^2+^) and potassium (K^+^), and chloride (Cl^-^) follow a circadian rhythm for a variety of plant tissues, which in turn regulate turgor changes that result in circumnutation of tendrils and folding-unfolding of leaves for several species (Scorza and Dornelas 2011; Stolarz 2009). Still, it remains unclear if the purpose of circumnutation and coiling movements in *C. campestris*; are influenced by its circadian clock and if these movements serve as a strategic tactic for successful parasitism. Further experiments are required to fully understand this biological aspect of *Cuscuta*.

Previous studies have shown that *Cuscuta* has loose or tight coils under different wavelengths, critically affecting its haustoria development (Furuhashi, et al. 2021; Pan, et al. 2022; Yokoyama, et al. 2023). When *Cuscuta* presents loose coils under red light or a high red/far red (R/FR) ratio, there was no haustoria development, while low R/FR ratio and blue lights induced tight coils and successful haustoria development. An interesting discovery from previous studies, although not directly focused on this aspect, is that the final coil at the upper section of loose coils consistently exhibited higher angles compared to tight coils, even though *Cuscuta* wraps around the host multiple times, with all coils varying in angles. Notably, haustoria organs were produced from the flattened upper coil of tight coils, indicating that the angle of the top coil is important for the haustoria development (Yokoyama, et al. 2023). In this study, we separately analyzed Coil 1, Coil 2, and stabilization of coiling (positioning and tightening) mathematically. As a result, there were significant angle and time differences when comparing Coil 1 and Coil 2. On one hand, Coil 1 tended to position for 5.7 hours and stabilize after 9.9 hours, coming to a final angle of 25° with respect to the base. On the other hand, Coil 2 tended to position just for 3.0 hours and stabilize after 6.0 hours, coming to a final angle of 10° with respect to the base. Therefore, Coil 1 tended to exhibit much more mobility after contacting the skewer to then stay at a slanted angle, whereas Coil 2 tended to be more limited in movement and settled at a flat angle (Supp Figs 2c, 3c). Thus, we suggest that our approach has capability to distinguish mechanistic differences between loose or tight coils, as previously described (Yokoyama, et al. 2023), and predict the development of the haustoria organ prior to its emergence from the inoculated *Cuscuta* stem.

Moreover, the automated pipeline of this study can determine more nuanced time points and periods related to *Cuscuta*’s coiling process. For instance, through these automatic analyses, it was observed that the time it takes *C. campestris* to twist around the skewer is independent of the time it takes to stabilize its coiling, regardless of the inoculation time. This suggests that twisting and stabilization follow two different mechanisms. This agrees with prior reports where twining and haustoria development could be independently induced from each other using different light stimuli (Furuhashi, et al. 2021). Another observation from automated annotations is that once *Cuscuta* initiates coiling, the period duration for each of its coiling stages —positioning, tightening, and stabilization— is independent of the time of inoculation, even though *Cuscuta* inoculated during the morning initiates considerably quicker than the one inoculated during afternoon. It was also observed that all phenotypes related to final coil positions and angles were also independent of inoculation time (Supp Fig 4).

At the moment, the proposed pipeline can only extract phenotypes associated to *C. campestris*’s stem angle and position with respect to the skewer by contrasting the skewers and parasites’ color. A more careful choice of background and skewer color can provide clearer images from which the whole *C. campestris* stem could be digitally isolated. Recent transgenic RUBY *C. campestris* exhibit robust betalain pigment expression, which would greatly facilitate its digital separation from the skewer and the background (Adhikari, et al. 2024). This setup can potentially solve our current shortcomings, and it would be thus possible to automatically extract more movement and circumnutation characteristics, and even refine the criteria to determine coil initiation and completion so that they coincide with those used in a manual setting.

Circumnutation and coiling represent crucial behaviors for stem parasitic plants, facilitating their effective parasitic lifestyle. This research endeavors to advance computational image analysis tools capable of monitoring *Cuscuta* movement under controlled laboratory conditions within a short period of time. Through these invested tools, we aim to delve into the biological, morphological, biochemical, and physiological aspects of *Cuscuta* mobility. Such investigations promise fundamental insights into the intricate interactions between plants, particularly between hosts and parasitic species, thereby laying the groundwork for parasitic plant management strategies.

In summary, our study introduces a Python-based image processing pipeline designed to phenotype key traits associated with *Cuscuta* circumnutation and coiling. This tool automates the detection of plant movement parameters such as coiling location, angle, initiation, and completion times, as well as the durations of various coiling stages. We utilized this pipeline to better understand the coiling behavior and circadian rhythms of *Cuscuta*. The data generated by this automated tool agreed with manual annotations, underscoring its accuracy and reliability.

Additionally, we have made the pipeline publicly available in the form of Jupyter notebooks on GitHub. This facilitates adaptations to diverse imaging datasets and experimental conditions, broadening its applicability. Our approach not only refines the accuracy of tracking plant movements but also lays the groundwork for further exploration into the mechanisms of parasitic plant interactions, potentially leading to sustainable methods for managing parasitic weeds.

## AUTHOR CONTRIBUTIONS

SP, EA, and JB conceived the ideas and designed the methodology. MB and SA conducted lab work and analyzed the data. EA performed image analysis and script development. MB and EA led the writing of the manuscript. All authors contributed critically to the drafts and gave final approval for publication.

## ACKNOWLEDGMENTS

This research received support by the Pests and Beneficial Species in Agricultural Production Systems, project award no. 2023-67013-39896, from the U.S. Department of Agriculture’s National Institute of Food and Agriculture to SP, as well as funding from the Research Council, the College of Agriculture, Food, and Natural Resources (CAFNR), and the Interdisciplinary Plant Group (IPG) at the University of Missouri. Grammarly generated responses to the following AI prompts: Prompts created by Grammarly- “Improve it”.

## CONFLICTING INTEREST STATEMENT

All authors declare that they have no conflicts of interest.

## DATA AVAILABILITY STATEMENT

The developed Python-based pipeline is available as a collection of Jupyter notebooks at https://github.com/ejamezquita/cuscuta/.

The video datasets used and/or analyzed during the current study are available here: https://youtube.com/playlist?list=PLZkYcVyQr2u4tT0yoZAkrMqzQxRIDvxru&feature=shared

## Notes

### Competing Interest Statement

The authors have declared no competing interest.

### Summary of Updates

Figure 3 has been updated, and the introduction, methods, result, and discussion part have been revised.

## REFERENCES

1. Adhikari S, Mudalige A, Phillips L, Lee H, Bernal-Galeano V, Gruszewski H, Westwood JH, Park S-Y (2024) Agrobacterium-mediated Cuscuta campestris transformation as a tool for understanding plant-plant interactions. bioRxiv 2024.2002.2023.581736

2. Agostinelli D, DeSimone A, Noselli G (2021) Nutations in plant shoots: endogenous and exogenous factors in the presence of mechanical deformations. Frontiers in Plant Science 12:608005

3. Araus JL, Cairns JE (2014) Field high-throughput phenotyping: the new crop breeding frontier. Trends Plant Sci 19:52–61

4. Ashton FM, Hutchison JM (1980) Germination of Field Dodder (Cuscuta campestris). Weed Sci 28:330–333

5. Baillaud L (1962) Handbuch der Pflanzenphysiologie. In. Berlin—Göttingen—Heidelberg Springer

6. Bernal-Galeano V, Beard K, Westwood JH (2022) An artificial host system enables the obligate parasite Cuscuta campestris to grow and reproduce in vitro. Plant Physiol 189:687–702

7. Brown AH (1993) Circumnutations: from Darwin to space flights. Plant Physiol 101:345

8. Cai C, Li J, Ren Z, Wan J, Xiao J, van Kleunen M (2022) Implications of climate change for environmental niche overlap between five Cuscuta pest species and their two main Leguminosae host crop species. Weed Sci 70:543-552

9. Creux N, Harmer S (2019) Circadian Rhythms in Plants. Cold Spring Harb Perspect Biol 11:

10. Darwin C, Darwin F (1880) The power of movement in plants. John Murray

11. Fahlgren N, Gehan M, Baxter I (2015) Lights, camera, action: high-throughput plant phenotyping is ready for a close-up. Curr Opin Plant Biol 24:93–99

12. Fischler MA, Bolles RC (1981) Random sample consensus: a paradigm for model fitting with applications to image analysis and automated cartography. Commun ACM 24:381–395

13. Furuhashi K, Iwase K, Furuhashi T (2021) Role of Light and Plant Hormones in Stem Parasitic Plant (Cuscuta and Cassytha) Twining and Haustoria Induction. Photochem Photobiol 97:1054–1062

14. Furuhashi T, Furuhashi K, Weckwerth W (2011) The parasitic mechanism of the holostemparasitic plant Cuscuta. J Plant Interactions 6:207–219

15. Goldwasser Y, Lanini WT, Sazo MRM (2012) Control of Field Dodder (Cuscuta campestris) Parasitizing Tomato with ALS-Inhibiting Herbicides. Weed Technol 26:740–746

16. Greenham K, McClung CR (2015) Integrating circadian dynamics with physiological processes in plants. Nature Reviews Genetics 16:598–610

17. Hartenstein M, Albert M, Krause K (2023) The plant vampire diaries: a historic perspective on Cuscuta research. J Exp Bot 74:2944–2955

18. Hegenauer V, Welz M, Körner M, Albert M (2017) Growth Assay for the Stem Parasitic Plants of the Genus Cuscuta. Bio Protoc 7:e2243

19. Jhu M-Y, Sinha NR (2022) Cuscuta species: Model organisms for haustorium development in stem holoparasitic plants. Frontiers in plant science 13:1086384

20. Kim G, Westwood JH (2015) Macromolecule exchange in *Cuscuta*–host plant interactions. Curr Opin Plant Biol 26:20–25

21. Kogan MJ, Lanini WT (2005) Biology and management of Cuscuta in crops. Ciencia E Investigacion Agraria 32:165–180

22. Mann HB, Whitney DR (1947) On a test of whether one of two random variables is stochastically larger than the other. The annals of mathematical statistics 50–60

23. Mao Y, Liu H, Wang Y, Brenner ED (2023) A deep learning approach to track Arabidopsis seedlings’ circumnutation from time-lapse videos. Plant Methods 19:18

24. Masanga J, Mwangi BN, Kibet W, Sagero P, Wamalwa M, Oduor R, Ngugi M, Alakonya A, Ojola P, Bellis ES, Runo S (2021) Physiological and ecological warnings that dodders pose an exigent threat to farmlands in Eastern Africa. Plant Physiol 185:1457–1467

25. McClung CR (2006) Plant circadian rhythms. Plant Cell 18:792–803

26. Millet B, Melin D, Bonnet B, Ibrahim C, Mercier J (1984) Rhythmic circumnutation movement of the shoots in Phaseolus vulgaris L. Chronobiol Int 1:11–19

27. Moulton DE, Oliveri H, Goriely A (2020) Multiscale integration of environmental stimuli in plant tropism produces complex behaviors. Proceedings of the National Academy of Sciences 117:32226–32237

28. Nadler-Hassar T, Rubin B (2003) Natural tolerance of Cuscuta campestris to herbicides inhibiting amino acid biosynthesis. Weed Res 43:341–347

29. Niinuma K, Nakagawa M, Calvino M, Mizoguchi T (2007) Dance of plants with circadian clock. Plant Biotechnol 24:87–97

30. Niinuma K, Someya N, Kimura M, Yamaguchi I, Hamamoto H (2005) Circadian Rhythm of Circumnutation in Inflorescence Stems of Arabidopsis. Plant and Cell Physiology 46:1423–1427

31. Pan H, Li Y, Chen L, Li J (2022) Molecular Processes of Dodder Haustorium Formation on Host Plant under Low Red/Far Red (R/FR) Irradiation. International Journal of Molecular Sciences 23:7528

32. Pierz LD, Heslinga DR, Buell CR, Haus MJ (2023) An image-based technique for automated root disease severity assessment using PlantCV. Applications in Plant Sciences 11:e11507

33. Riviere S, Clayson C, Dockstader K, Wright MAR, Costea M (2013) To attract or to repel? Diversity, evolution and role of the “most peculiar organ” in the Cuscuta flower (dodder, Convolvulaceae)—the infrastaminal scales. Plant Syst Evol 299:529–552

34. Robil JM, Gao K, Neighbors CM, Boeding M, Carland FM, Bunyak F, McSteen P (2021) grasviq: an image analysis framework for automatically quantifying vein number and morphology in grass leaves. The Plant Journal 107:629–648

35. Runyon JB, Mescher MC, De Moraes CM (2006) Volatile chemical cues guide host location and host selection by parasitic plants. Science 313:1964–1967

36. Saric-Krsmanovic M, Vrbnicanin S (2015) Field dodder - how to control it? Pestic Fitomed 30:137–145

37. Savitzky A, Golay MJ (1964) Smoothing and differentiation of data by simplified least squares procedures. Analytical chemistry 36:1627–1639

38. Scorza LCT, Dornelas MC (2011) Plants on the move: Towards common mechanisms governing mechanically-induced plant movements. Plant Signaling & Behavior 6:1979–1986

39. Shimizu K, Aoki K (2019) Development of Parasitic Organs of a Stem Holoparasitic Plant in Genus Cuscuta. Frontiers in Plant Science 10:

40. Stolarz M (2009) Circumnutation as a visible plant action and reaction: physiological, cellular and molecular basis for circumnutations. Plant Signal Behav 4:380–387

41. Sun G, Xu Y, Liu H, Sun T, Zhang J, Hettenhausen C, Shen G, Qi J, Qin Y, Li J, Wang L, Chang W, Guo Z, Baldwin IT, Wu J (2018) Large-scale gene losses underlie the genome evolution of parasitic plant Cuscuta australis. Nature Communications 9:2683

42. Swartz LG, Liu S, Dahlquist D, Kramer ST, Walter ES, McInturf SA, Bucksch A, Mendoza- Cózatl DG (2023) OPEN leaf: an open-source cloud-based phenotyping system for tracking dynamic changes at leaf-specific resolution in Arabidopsis. The Plant Journal 116:1600–1616

43. Těšitel J (2016) Functional biology of parasitic plants: a review. Plant Ecology and Evolution 149:5–20

44. Wu Y, Xie J, Wang L, Zheng H (2020) Circumnutation and Growth of Inflorescence Stems of Arabidopsis thaliana in Response to Microgravity under Different Photoperiod Conditions. Life 10:26

45. Yokoyama T, Watanabe A, Asaoka M, Nishitani K (2023) Germinating seedlings and mature shoots of Cuscuta campestris respond differently to light stimuli during parasitism but not during circumnutation. Plant, Cell Environ 46:1774–1784

46. Yoshihara T, Iino M (2005) Circumnutation of rice coleoptiles: its occurrence, regulation by phytochrome, and relationship with gravitropism. Plant, Cell Environ 28:134–146

47. Zhang TY, Suen CY (1984) A fast parallel algorithm for thinning digital patterns. Communications of the ACM 27:236–239

