## Supplementary material for "The Early Dodder Gets the Host: Decoding the Coiling Patterns of *Cuscuta campestris* with Automated Image Processing": Supp Figures and Tables

SUPPLEMENTARY MATERIALS

**Supplementary Fig. 1**. The experiment setting of *Cuscuta* inoculation and the time-lapse. Far-red enriched lights were installed for successful coiling on live (*Arabidopsis*) and non-live (bamboo skewer) hosts.

**Supplementary Fig. 2.** Distance between Coil 1 and Coil 2 made by *Cuscuta* stems around the bamboo skewers at three different inoculation times (9 AM, 12 PM, and 4 PM) **(a)**, time for *Cuscuta* to complete a 360º twist **(b)**, angles of *Cuscuta* coils with respect to the base in degrees **(c)**, and position height above tapes **(d)**. ns: Not significant, p-value of ≥ 0.05; *: p-value of 0.005 to 0.05, **: p-value of 0.0005 to 0.005, and ***: p-value < 0.0005.

**Supplementary Fig. 3.** Time period duration of each coil positioning (a). Time period duration of each coil tightening (b). Time for each coil to stabilize once it has initiated (c) Total time spent coiling **(d).** ns: Not significant, p-value of ≥ 0.05; *: p-value of 0.005 to 0.05, **: p-value of 0.0005 to 0.005, and ***: p-value < 0.0005.

**Supplementary Fig. 4.** Comparison of different time period durations of different stages of coiling, depending on the Coil and the inoculation time. $r_s$ indicates the Spearman coefficient of correlation. ns: Not significant, p-value of ≥ 0.05; *: p-value of 0.005 to 0.05, **: p-value of 0.0005 to 0.005, and ***: p-value < 0.0005.

**Supplemental Video 1.** Time-lapse video of *Cuscuta* coiling on non-live and live hosts.

**Supplemental Video 2.** Representative time-lapse video of *Cuscuta* inoculated at 9 AM.

**Supplemental Video 3.** Representative time-lapse video of *Cuscuta* inoculated at 12 PM.

**Supplemental Video 4.** Representative time-lapse video of *Cuscuta* inoculated at 4 PM.

**Supplemental Video 5.** Automated image analysis of *Cuscuta* coiling corresponding to Figure 4. (Top left) Section of the original photo. Distance to the point of inoculation (tape) indicated in millimeters in the y-axis. (Bottom left) Same section but the skewer's center (red), *Cuscuta*'s skeleton (yellow), crossing points (white stars), Coil 1 angle (light blue), and Coil 2 angle (mustard) are overlaid. (Top right) Smoothened time series of Coil's angle and (bottom right) Coil's distance to point of inoculation. Black bar at the bottom indicates the passage of time.

**Supplemental Videos 6 through 9**. Similar analysis as Suppl. Video 5 for the rest of skewers in the repetition.

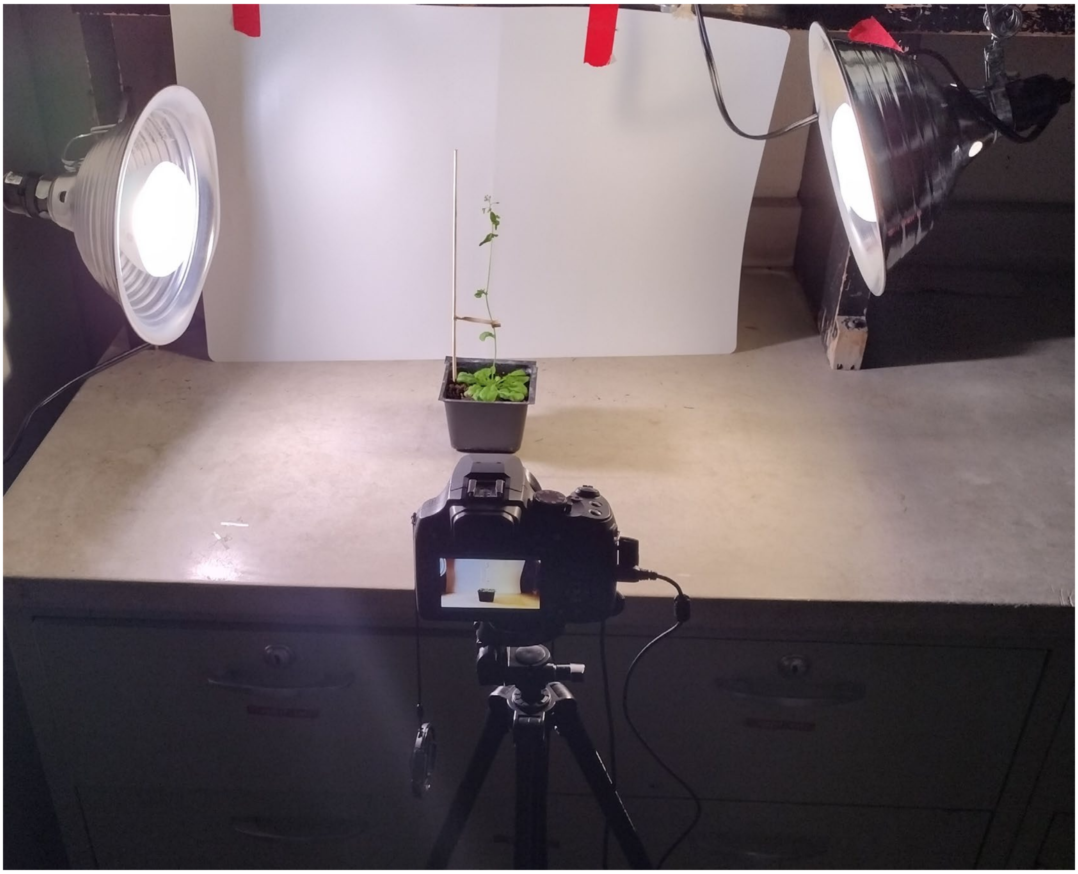

**Supplementary Fig. 1.**

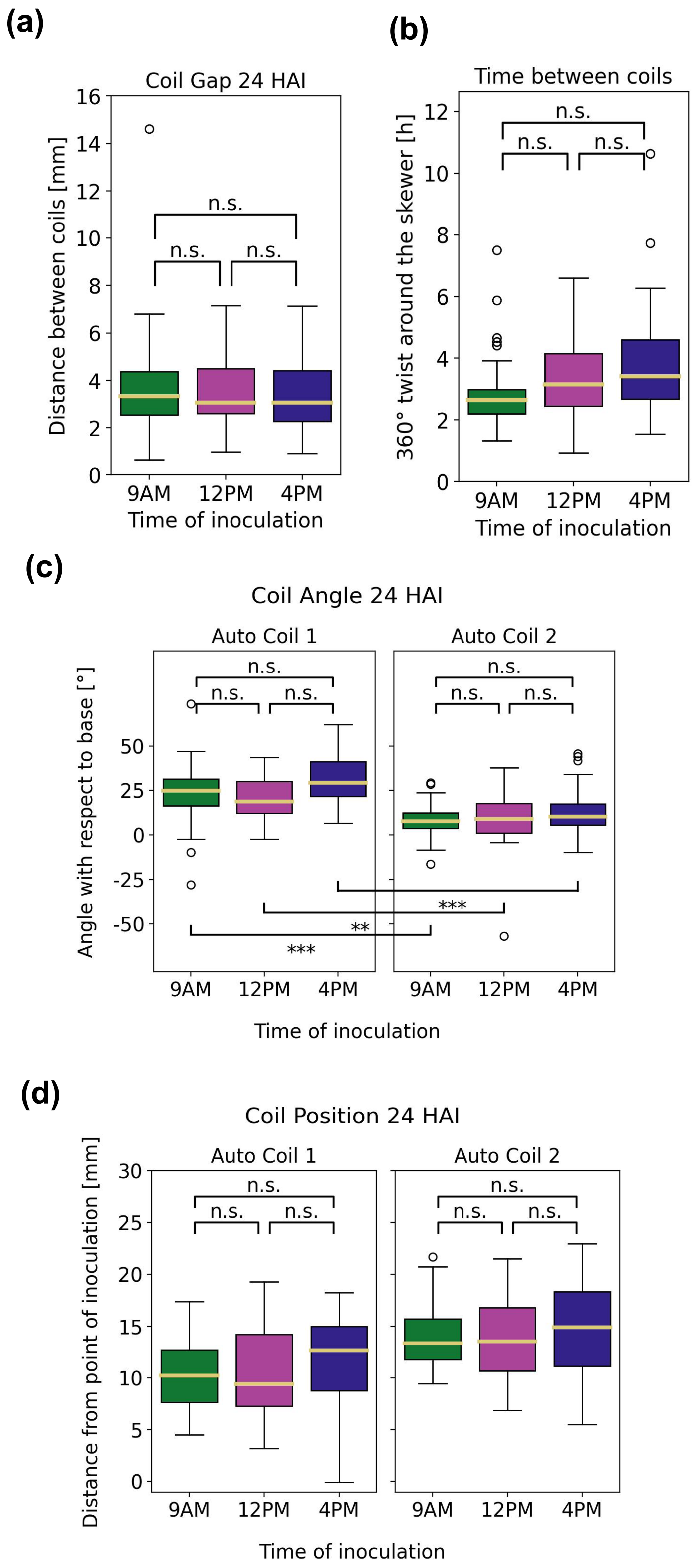


**Supplementary Fig. 2.**

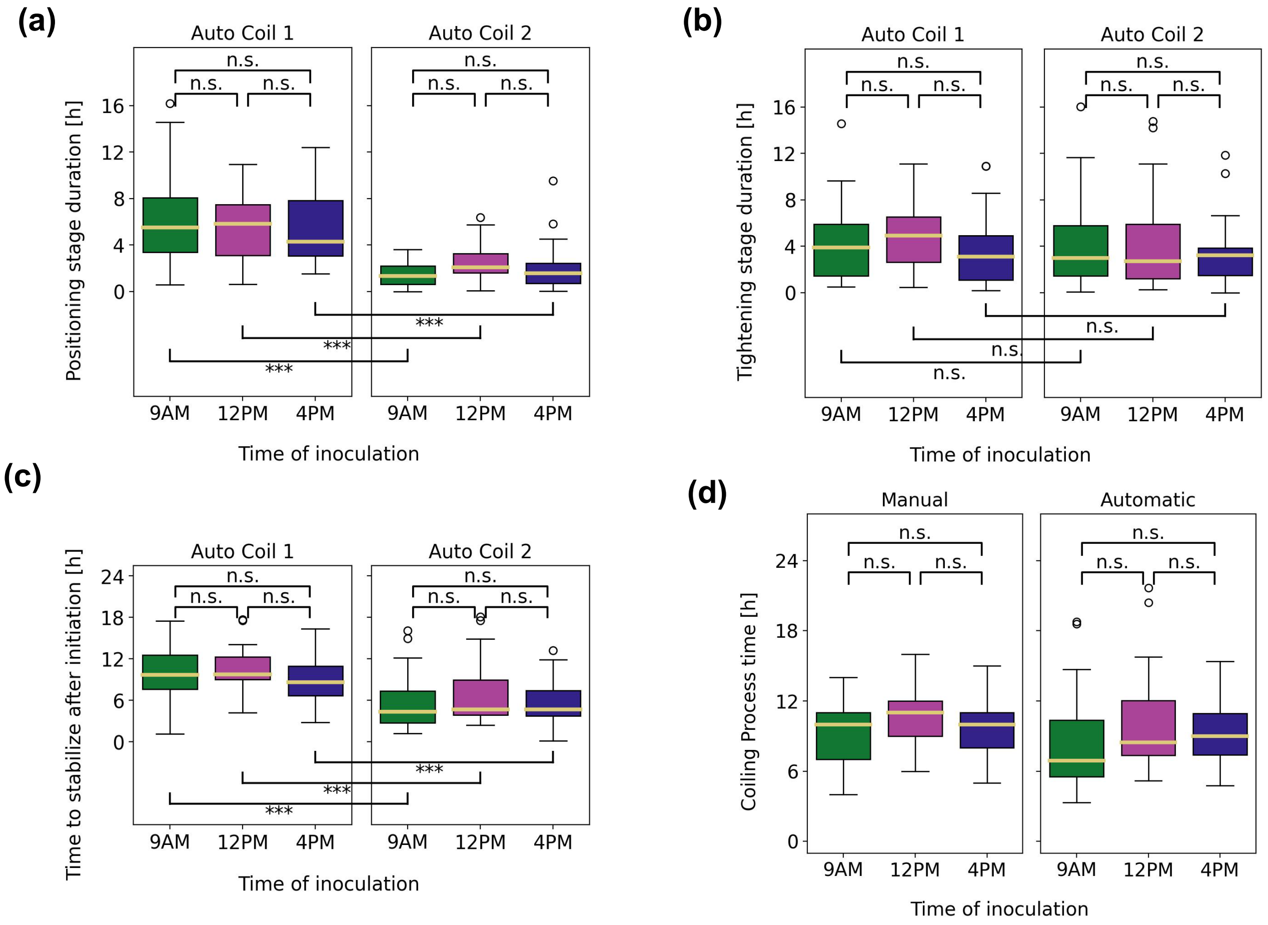


**Supplementary Fig. 3.**

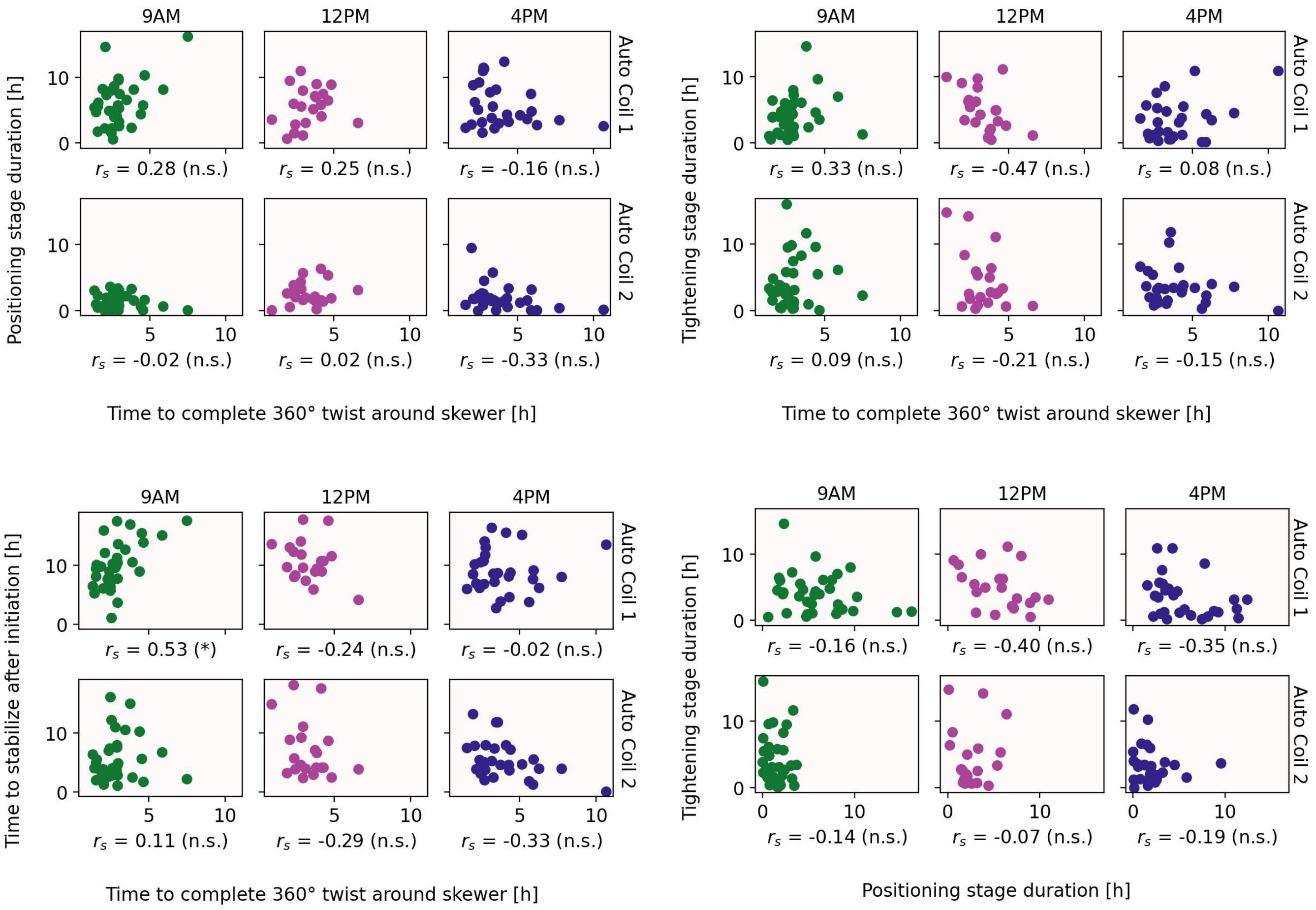


**Supplementary Fig. 4.**

**Supp table 1.** The success rates of each *Cuscuta* coil were counted by the manual observation method. Five *Cuscuta* stems were attached on five different bamboo skewers under three different inoculation times (9 AM, 12 PM, and 4 PM), showing the number of coiled *Cuscuta* per number of applied *Cuscuta* stems.

| | | Inoculation time | | |
|---|---|---|---|---|
| | | **9 AM** | **12 PM** | **4 PM** |
| **Replicates number** | Rep 1 | 5/5 | 4/5 | 4/5 |
| | Rep 2 | 5/5 | 4/5 | 4/5 |
| | Rep 3 | 5/5 | 3/5 | 5/5 |
| | Rep 4 | 5/5 | 4/5 | 4/5 |
| | Rep 5 | 5/5 | 5/5 | 3/5 |
| | Rep 6 | 5/5 | 4/5 | 4/5 |
| | Rep 7 | 5/5 | 5/5 | 5/5 |
| **Total number of successful *Cuscuta* coils** | | 35/35 | 29/35 | 29/35 |

**Supp table 2.** The success rates of each *Cuscuta* coil counted by the automated detection method. Five *Cuscuta* stems were attached on five different bamboo skewers under three different inoculation times (9 AM, 12 PM, and 4 PM), showing the number of coiled *Cuscuta* per number of applied *Cuscuta* stems.

| | | **Inoculation time** | | |
|---|---|---|---|---|
| | | **9 AM** | **12 PM** | **4 PM** |
| **Replicates number** | Rep 1 | 5/5 | 4/5 | 4/5 |
| | Rep 2 | 4/5 | 3/5 | 4/5 |
| | Rep 3 | 5/5 | 3/5 | 5/5 |
| | Rep 4 | 5/5 | 4/5 | 4/5 |
| | Rep 5 | 5/5 | 3/5 | 3/5 |
| | Rep 6 | 5/5 | 4/5 | 3/5 |
| | Rep 7 | 5/5 | 4/5 | 5/5 |
| **Total number of successful *Cuscuta* coils** | | 34/35 | 25/35 | 28/35 |

**Supp table 3**. Coiling initiation, completion, and coiling process time counted by the manual observation method. HAI: hours after inoculation. SEM: Standard error of the mean.

| | **Initiation time** | | **Completion time** | | **Coiling process** | |
|---|---|---|---|---|---|---|
| | **Mean (HAI)** | **SEM** | **Mean (HAI)** | **SEM** | **Mean (Hours)** | **SEM** |
| **9 AM** | 5.26 | 0.32 | 14.43 | 0.57 | 9.17 | 0.44 |
| **12 PM** | 5.90 | 0.50 | 16.48 | 0.55 | 10.59 | 0.43 |
| **4 PM** | 8.45 | 0.56 | 18.00 | 0.57 | 9.55 | 0.44 |

**Supp table 4.** Coiling initiation, completion, and coiling process time of the 1$^{st}$ coil counted by the automatic detection method. HAI: hours after inoculation. SEM: Standard error of the mean.

| | **Initiation time** | | **Completion time** | | **Coiling process** | |
|---|---|---|---|---|---|---|
| | **Mean (HAI)** | **SEM** | **Mean (HAI)** | **SEM** | **Mean (Hours)** | **SEM** |
| **9 AM** | 4.83 | 0.21 | 14.83 | 0.75 | 10.00 | 0.68 |
| **12 PM** | 5.29 | 0.33 | 15.88 | 0.81 | 10.59 | 0.72 |
| **4 PM** | 7.93 | 0.49 | 16.65 | 0.68 | 8.72 | 0.63 |

**Supp table 5.** Coiling initiation, completion, and coiling process time of the 2$^{nd}$ coil counted by the automatic detection method. HAI: hours after inoculation. SEM: Standard error of the mean.

| | **Initiation time** | | **Completion time** | | **Coiling process** | |
|---|---|---|---|---|---|---|
| | **Mean (HAI)** | **SEM** | **Mean (HAI)** | **SEM** | **Mean (Hours)** | **SEM** |
| **9 AM** | 7.76 | 0.32 | 13.32 | 0.71 | 5.56 | 0.63 |
| **12 PM** | 8.65 | 0.47 | 15.18 | 0.79 | 6.53 | 0.88 |
| **4 PM** | 11.90 | 0.53 | 17.35 | 0.64 | 5.46 | 0.59 |
